# Defining antigen targets to dissect vaccinia virus (VACV) and Monkeypox virus (MPXV)-specific T cell responses in humans

**DOI:** 10.1101/2022.09.06.506534

**Authors:** Alba Grifoni, Yun Zhang, Alison Tarke, John Sidney, Paul Rubiro, Maria Reina-Campos, Gilberto Filaci, Jennifer Dan, Richard H. Scheuermann, Alessandro Sette

## Abstract

The current Monkeypox virus (MPXV) outbreak in non-endemic countries is raising concern about the pandemic potential of novel orthopoxviruses. Little is known regarding MPXV immunity in the context of MPXV infection or vaccination with Vaccinia-based vaccines (VACV). As with vaccinia, T cells are likely to provide an important contribution to overall immunity to MPXV. Here we leveraged the epitope information available in Immune Epitope Database (IEDB) on VACV to predict potential MPXV targets recognized by CD4^+^ and CD8^+^ T cell responses. We found a high degree of conservation between VACV epitopes and MPXV, and defined T cell immunodominant targets. These analyses enabled the design of peptide pools able to experimentally detect VACV-specific T cell responses and MPXV cross-reactive T cells in a cohort of vaccinated individuals. Our findings will facilitate the monitoring of cellular immunity following MPXV infection and vaccination.

## Introduction

An outbreak of Monkeypox virus (MPXV) has been reported in 96 countries. On August 24^th^, 2022, the WHO reported 41,664 confirmed cases of MPXV infection and five deaths in non-endemic regions with a prevalence of cases in the European region (50%) followed by the Americas (49%);(https://www.who.int/publications/m/item/multi-country-outbreak-of-monkeypox--external-situation-report--4---24-august-2022). While MPXV infections and outbreaks have been reported in the past three decades on the African continent, specifically the Democratic Republic of Congo, Nigeria and Central African Republic, this current outbreak is unprecedent in size and scope, having spread globally to almost 100 countries, the vast majority of which have not historically reported MPXV cases, including Europe and the U.S. (Bunge et al., 2022).

This alarming 2022 MPXV outbreak is being monitored by health and regulatory agencies worldwide and has received considerable attention in the general news. This awareness is particularly acute in the context of heightened consideration of potential global infectious disease outbreaks based on the SARS-CoV-2 pandemic experience. With the current outbreak, it is important to understand immunity against MPXV. However, only a few studies thus far have addressed immune responses to MPXV infections in humans (Hammarlund et al., 2008; Hammarlund et al., 2005; Karem et al., 2007), and essential knowledge gaps are apparent. First, little to no information is available regarding the quality and duration of immune responses associated with natural MPXV infection in humans. Second, very little actual efficacy data in humans is available for the MPXV vaccines, all based on the Vaccinia Virus (VACV), as discussed in more detail below. Third, it is unknown to what extent the cellular immune response induced in humans by VACV vaccination are cross-reactive with MPXV epitopes. Finally, the knowledge gaps described above should be addressed not only in relation to MPXV in general, but also in relation to the founder strain associated with the current outbreak, which may be associated with peculiarities in terms of transmissibility/infectivity, and responses induced by infection (Ahmed et al., 2022).

A large amount of information is available regarding immune responses and correlates of protection from VACV infection (Amanna et al., 2006; Kennedy et al., 2009; Xu et al., 2004), the virus utilized as a vaccine to protect from smallpox disease caused by variola virus (VARV) infection (Carroll and Moss, 1997). Several VACV studies demonstrate that antibody responses are crucial for prevention of disease (Amanna et al., 2006; Xu et al., 2004), whereas T cell responses are important to control and terminate Pox virus infections (Jing et al., 2005; Pauli et al., 2010). There is a significant body of information available in the peer-reviewed literature about T cell epitopes for orthopoxviruses (OPXV) in general, and VACV in particular (Drexler et al., 2003; Jing et al., 2005; Jing et al., 2008; Moutaftsi et al., 2006; Oseroff et al., 2005; Stone et al., 2005; Terajima et al., 2003; Terajima et al., 2008). However, only two studies have investigated T cell responses against MPXV in humans (Alzhanova et al., 2014; Song et al., 2013).

VACV was utilized (under the brand name of Dryvax) to eradicate smallpox in the 1980’s (Jacobson et al., 2009). However, Dryvax vaccinated individuals with impaired cellular immunity were found deficient in VACV viral control (Howell et al., 2006; Redfield et al., 1987), and at risk of severe reactions and other safety issues (Jacobs et al., 2009). Accordingly, Dryvax vaccination (and vaccination with the related attenuated vaccine Acambis 2000) was largely discontinued after 2001, and replaced by the MVA (Modified Vaccinia Ankara) virus (under the brand name JYNNEOS) which has a superior safety profile (Xiang and White, 2022), and induces similar levels of antibody responses (Davies et al., 2008).

Studies in non-human primate models using VACV vaccination to prevent MPXV infection demonstrate efficacy in preventing infection and/or attenuating disease severity (Earl et al., 2004; Edghill-Smith et al., 2005b; Karem et al., 2007; Nigam et al., 2007). Data from human studies is limited to a single observational study demonstrating 85% efficacy in preventing MPXV disease in subjects vaccinated with Dryvax (Jezek et al., 1986). The Jynneos vaccine is approved for use to prevent MPXV infection/disease based on serological responses, but no data is available addressing clinical efficacy in humans. An additional knowledge gap is the degree of CD4^+^ and CD8^+^ T cell epitope conservation elicited by VACV vaccination for MPXV infection. MPXV shares 90% or more overall sequence homology with VACV and VACV-like cowpox viruses (Babkin et al., 2022), suggesting that VACV-induced T cell responses might be largely cross-reactive with MPXV epitope sequences. Indeed, we previously noted that VACV epitopes are largely conserved in VARV (Sette et al., 2009), thus suggesting a similar strategy can be applied to other orthopoxviruses. Here, we used the Immune Epitope Database and Analysis Resource (IEDB) (Vita et al., 2019) and Virus Pathogen Resource (ViPR) (Pickett et al., 2012) to compile known epitope information on orthopoxviruses in general, and VACV in particular, to determine protein conservation, and assess immunodominance for both CD4^+^ and CD8^+^ T cell responses.

Previous work from our group has shown that pools of a large number of peptides (megapools; MP) (Carrasco Pro et al., 2015) generated by sequential lyophilization can be used to easily and accurately measure CD4^+^ and CD8^+^ T cell responses against complex antigens. These MP can be based on overlapping peptides spanning the entire sequence of an antigen of interest, or more frequently either experimentally defined or predicted HLA binders/T cell epitopes. This approach has been validated for a number of allergens, and bacterial and viral targets (Arlehamn et al., 2012; da Silva Antunes et al., 2017; Grifoni et al., 2019; Grifoni et al., 2020b; Grifoni et al., 2020c), but MPs for poxviruses have not been described as yet. In this study, we develop pools of previously defined epitopes that can be utilized to assess T cell responses to VACV. In parallel, we further apply validated bioinformatic tools to predict MPXV T cell epitopes likely to be recognized in humans based on MPXV ortholog proteins that are immunodominant in VACV responses. Finally, we validate the biological activity of these reagents by testing banked PBMCs from previously described (Oseroff et al., 2005) Dryvax vaccinated subjects.

## Results

### Orthopox T cell epitopes curated in the IEDB

In the first analysis, we inventoried the Orthopoxvirus data in the IEDB, as of the end of May, 2022. We considered data curated from the published literature, as well as data provided by direct database submission, and related to CD4^+^ (class II restricted) and CD8^+^ (class I restricted) T cell epitopes (see Methods section for further details). The analysis identified 318 unique CD4^+^ and 659 unique CD8^+^ T cell epitopes derived from orthopoxviruses (**Supplemental Table 1**). As expected, the vast majority of epitopes (88%) have been described in the context of VACV, with minor components represented by epitopes defined in the context of VAR and ectromelia viruses. The majority of CD4^+^ epitopes (78%) and slightly more than a third of the CD8 epitopes were associated with responses in humans (**Supplemental Table 1**).

Based on these data, we next developed two MPs, OPX-CD8-E and OPX-CD4-E, to probe responses to orthopoxviruses in human donors. Specifically, for OPX-CD8-E we selected the 238 CD8^+^ epitopes recognized in humans, while for OPX-CD4-E we included all 318 CD4^+^ epitopes recognized in any species, based on the general high degree of overlap between binding repertoires of MHC class II molecules (Peters et al., 2020). As several of the reported CD4^+^ epitopes largely overlap or nest other epitopes, we performed a clustering analysis to create epitope regions of up to 22 residues that encompass nested or largely overlapping epitopes. Accordingly, a set of 300 CD4^+^ epitopes was generated. The CD4^+^ and CD8^+^ epitopes included in the respective pools are listed in **Supplemental Table 1**.

### Conservation of Orthopox epitopes within MPX

The MPXV and VACV viruses have been reported (Babkin et al., 2022) to share a high degree of sequence homology and conservation, suggesting that the T cell epitopes previously identified in other orthopoxviruses (mostly in VACV) may be conserved in MPX (Ahmed et al., 2022). Here, the MPXV_USA_2022_MA001 (MA001) isolate (GenBank accession ON563414) was selected as the representative strain of the 2022 MPXV outbreak because MA001 was the first sequence from the 2022 outbreak deposited in GenBank (Gigante et al., 2022) and had been annotated by BV-BRC (www.bv-brc.org) for ORF and protein sequences.

To identify the MPXV homologs of VACV antigens, a BLAST based homology search was used. All the 26 VACV antigens related to IEDB T cell epitopes except for VP8 and A47 had high identity (>83%) and good length coverage in MA001. With regard to the VP8 antigen, an alignment of MA001 with the RefSeq Zaire strain showed that MA001 had a 1 nt insertion in the VP8 region which caused a frameshift. This was likely a sequencing artifact. Therefore, this 1 nt insertion was removed, and the genome sequence was re-annotated using the GATU tool in ViPR (www.viprbrc.org) to obtain the VP8 homolog in MPXV. For the A47 antigen, neither MA001 nor the RefSeq Zaire strain had a good match. This antigen is unlikely to have a homolog in MPXV.

To assess the sequence conservation of the OPX-CD8-E and OPX-CD4-E epitope pools in MPXV, a k-mismatch string search program was developed to find all matched sequences of an input epitope. The matched sequences meet the criteria of having the same length as the input epitope, and having at most k mismatched residues (including terminal indels if there are any) in comparison to the input epitope. The terminal indels were allowed in order to accommodate the scenario of having indels at the beginning or end of a protein. In case of multiple matches for the same input epitope, the program also picks the optimal match.

The results indicate that both CD4^+^ and CD8^+^ epitopes are highly conserved with 94% of the CD4^+^ and 82% of the CD8^+^ epitopes being 100% conserved in MPXV (**Supplemental Table 1**), with high conservation (range 74% to 96%) irrespective of the viral species in which the epitopes were originally defined and described. Overall, the majority of the orthopox T cell epitopes included in OPX-CD8-E and OPX-CD4-E should potentially be useful to monitor responses to vaccination, and because of their conservation in MPXV, also potentially be useful to monitor T cell responses in the context of MPXV infection. Next, we validated the use of these pools to detect responses from VACV vaccinated subjects.

### T cell immune responses after Dryvax vaccination are detected by the Orthopox MPs

We evaluated T cell responses from cryopreserved PBMCs from a cohort of donors vaccinated with Dryvax, and a control cohort of non-vaccinated subjects, for their capacity to recognize the Orthopox MPs described above. The PBMCs from the vaccinated cohort are cross-sectional samples collected in 2005 (Oseroff et al., 2005) from healthcare workers who received Dryvax vaccination as a prophylactic measure against laboratory exposure of VACV. In most cases, the donors had previously been immunized in the context of childhood vaccination and/or previous occupational vaccination, and therefore the Dryvax vaccination would be a secondary (booster) immunization. The unexposed cohort encompassed donors born after 1980, to exclude previous childhood vaccination, and with no history of occupational vaccination or exposure. General characteristics of the cohorts are provided in **Supplemental Table 2**, and an in-depth description is provided in the method section.

To measure the T cell responses to OPX-CD8-E and OPX-CD4-E, we combined activation-induced marker (AIM) assays with cytokine intracellular staining (ICS) as recently described (Tarke et al., 2022). CD4^+^ and CD8^+^ T cell responses were measured by AIM as OX40^+^CD137^+^CD4^+^ T cells or CD69^+^CD137^+^CD8^+^ T cells following the gating strategy in **Figure S1A**. In the case of CD4^+^ T cells in the AIM assay, the highest responses were observed in recently vaccinated subjects, as expected. The magnitude of T cell reactivity was low and comparable pre-vaccination and 5-7 months post vaccination (GM±GSD; pre=0.09±2.90, 2 weeks=0.26±2.53, 5-7 months=0.11±2.86; P=0.004 Kruskal-Wallis test; Mann-Whitney comparisons in **Figure 1A**). The frequency of CD4^+^ T cell responders significantly increased from 52% to 95% 2 weeks post-vaccination with a decline to 61% 5-7 months post-vaccination (χ^2^ p= 0.01).

**Figure 1.**
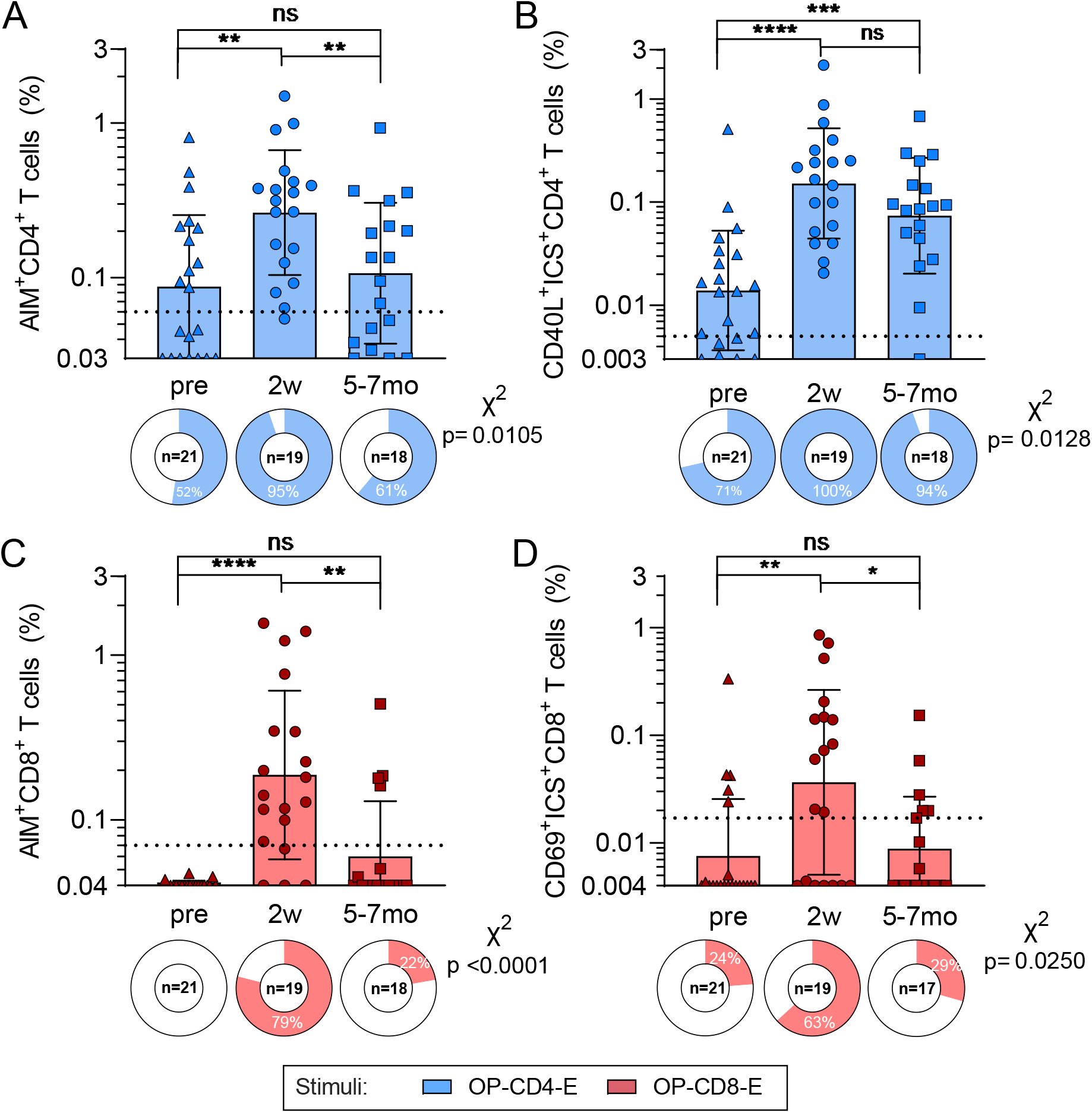
T cell responses against Orthopox MPs after Vaccinia vaccination. Refers to Figure S1-S2 and Supplemental Tables 1 to 4. PBMCs from each time-point were tested in AIM/ICS assays with experimentally defined OPXV MPs, OP-CD4-E (blue) and OP-CD8-E (red). OP-CD4-E specific CD4^+^ T cell reactivity are shown for (**A**) AIM and (**B**) ICS. OP-CD8-E specific CD8^+^ T cell reactivity are shown for (**C**) AIM and (**D**) ICS. The Y axis of each bar graph starts at the LOD and the LOS is indicated with a dotted line. Pie charts below each bar indicate the frequency of positive responders. Mann-Whitney T test was applied to each graph and p values symbols are shown when significant. * p<0.05, **p<0.01, ***p<0.001, ****p<0.0001. X^2^ test was applied to the frequencies of positivity and p values are listed on the right of each graph.

Similar findings were observed in the ICS readout, although CD4^+^ T cell responses showed a more modest and insignificant decline 5-7 months post-vaccination; responses post-vaccination were significantly higher in magnitude and frequency than those observed pre-vaccination (GM±GSD; pre=0.01±3.80, 2 weeks=0.15±3.42, 5-7 months=0.07±3.64; P<0.0001 Kruskal-Wallis test, χ^2^ p=0.0128, Mann-Whitney comparisons in **Figure 1B**). Non-vaccinated (unexposed) subjects did not yield appreciable responses, suggesting that the frequency of background exposure and/or pre-existing cross-reactive responses is low, at least in the cohort investigated (**Figure S1B-C**). The ICS assays demonstrated that CD4^+^ T cell responses were Th1 or cytotoxic responses, encompassing mostly granzyme B (GZMB) and IFNγ, followed by TNFα and IL-2 responses, in combination with CD40L expression (**Figure S2**). Significant increases in TNFα and IL-2 and decrease of GZMB production were observed in the quality of the T cell responses at 5-7 months post-vaccination, as compared to 2 weeks (TNFα: 2 weeks= 18% 5-7 months= 33% p=0.0088; IL-2: 2 weeks= 3% 5-7 months= 13% p=0.0059; GZMB: 2 weeks= 53% 5-7 months= 35% p=0.0292). A prevalence of CD40L^+^IFNγ^+^GZMB^+^ population followed by CD40L^+^GZMB^+^, CD40L^+^IFNγ^+^ and CD40L^+^TNFα^+^ populations were observed (**Figure S2A and S2E**).

In the case of CD8^+^ T cells for AIM assays, as expected the highest responses were also observed in recently vaccinated subjects, with the 5-7 months post-vaccination time point reverting to a magnitude of T cell reactivity similar to that observed pre-vaccination (GM±GSD; pre=0.04±1.04, 2 weeks=0.19±3.24, 5-7 months=0.06±2.15; P<0.0001 Kruskal-Wallis test; Mann-Whitney comparisons in **Figure 1C**). The frequency of positive CD8^+^ T cell responses increased to 79% 2 weeks post immunization and decreased to 22% at the 5-7 months’ time point (χ^2^ p<0.0001). Similar results were observed in the case of the ICS assay, where IFNγ, GZMB, TNFα, and IL-2 secreting CD69^+^CD8^+^ T cells were quantified (GM±GSD; pre=0.01±3.39, 2 weeks=0.04±7.22, 5-7 months=0.01±3.06; P=0.01 Kruskal-Wallis test; χ^2^ p=0.025; Mann-Whitney comparisons in **Figure 1D**). CD8^+^ T cell functionality was also assessed based on IFNγ, TNFα, IL-2, and/or GZMB expression and showed a prevalence of IFNγ followed by IL-2 and GZMB (**Figure S2C-F**).

### Definition of immunodominant ORFs

Pox viruses have relatively large genomes, encoding approximately 200 different ORFs that are expressed at different time points during the viral life cycle (Babkin et al., 2022). Several studies report broad and diverse immune responses to many different ORFs (Moutaftsi et al., 2006). To operationally define immunodominant Orthopoxvirus proteins, we assessed the distribution of the IEDB epitopes as a function of the corresponding source proteins (**Supplemental Table 1** and **Supplemental Table 3**). Accordingly, we identified 19 and 40 antigens associated with five or more reported CD4^+^ and CD8^+^ epitopes. The 19 immunodominant CD4^+^ ORFs accounted for 67% of the IEDB epitopes, and the 40 immunodominant CD8^+^ ORFs accounted for 61% of the IEDB epitopes **(Figure 2**).

**Figure 2.**
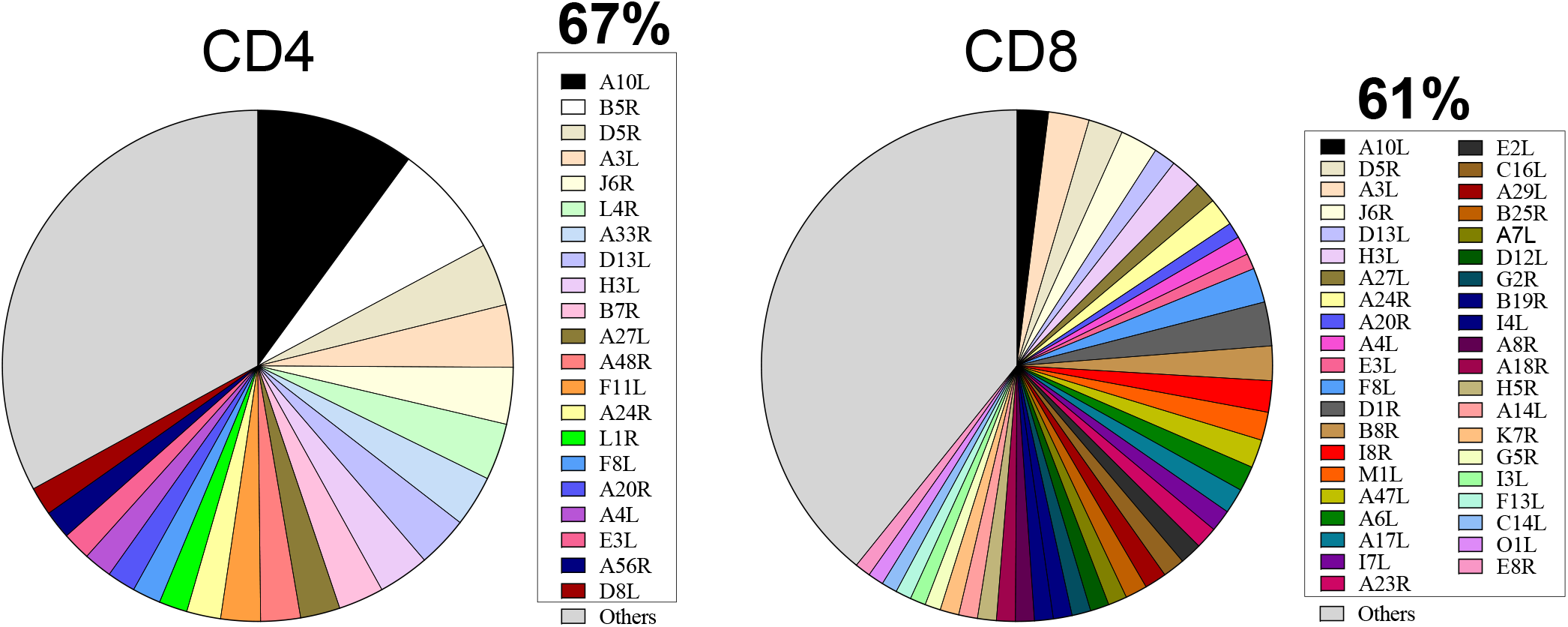
Orthopox protein immunodominance. Refers to Supplemental Table S1. The IEDB was mined for experimentally defined epitopes derived from Orthopoxviruses. A total of 47 different antigens were identified as the protein source for defined epitopes, corresponding to 19 and 39 antigens associated with CD4^+^ or CD8^+^ epitopes respectively. The pie charts represent the total proteins recognized by CD4^+^ and CD8^+^ as defined by the IEDB and listed in Supplemental Table S1. The most immunodominant antigens are listed in the figure legend and the remaining antigens (“others”) are colored in grey.

In general, immunodominant targets recognized by T cells identified by the analysis were mostly early transcribed proteins (Moss, 2011). The analysis further identified several targets, such as A10L, A3L, D5R, A26L/A30L, H3L, J6R, A27L, D13L, A24R, A4L and F8L, which are of particular interest, since they were dominant (as defined above) for both CD4^+^ and CD8^+^ T cells. The study of immunodominant ORFs is also of particular relevance considering the Jynneos vaccine is based on MVA, an attenuated VACV that has lost 14% of the original genome (Volz and Sutter, 2017), retaining only 157 ORFs (GenBank: U94848.1) (Antoine et al., 1998). Importantly, all the ORFs considered as dominant for either CD4^+^, CD8^+^ or both were also conserved in the MVA/Jynneos sequences. Taken together, these results confirm the extremely large breadth of immunogenic ORFs in Orthopox viruses (Oseroff et al., 2005). Simultaneously, these results demonstrate that a relatively manageable number of ORF accounting for the majority of responses can be defined.

### Prediction of MPX T cell epitopes from immunodominant ORFs

We next focused on the selected immunodominant ORFs to specifically predict MPXV epitopes, and design megapools expected to be suitable for detecting T cell responses in MPXV infected subjects. Accordingly, we predicted potential T cell epitopes from the MPXV orthologs of the 19 CD4^+^ and 40 CD8^+^ immunodominant antigens using tools provided in the IEDB analysis resource (Dhanda et al., 2019). For CD4^+^, we predicted 276 promiscuous HLA class II binders, following the method described by (Paul et al., 2015) (see further details in the Methods). In parallel, we also predicted a total of 1647 potential CD8^+^ epitopes binding to a panel of common HLA class I alleles. This approach was previously utilized in other viral systems (Grifoni et al., 2020a; Grifoni et al., 2020b) (see further details in the Methods).

In conclusion, the analysis described above identified 1923 predicted HLA binders derived from MPXV sequences. **Supplemental Table 3** provides a summary of the predicted CD4^+^ and CD8^+^ T cell epitopes as a function of the source protein. These epitopes were utilized to develop specific MPs potentially useful to monitor cross-reactive responses to vaccination, and MPXV-specific responses after infection. Specifically, we designed one MP specific for CD4^+^ responses (MPXV-CD4-P), while for CD8^+^ we pooled the predicted peptides as a function of the protein of origin into 5 total pools (MPXV-CD8-P1-P5) (**Supplemental Table 3** and **Supplemental Table 4**).

### Assessment of CD4^+^ and CD8^+^ T cell cross-reactive responses able to recognize MPXV in smallpox vaccinated individuals

We then evaluated whether VACV-induced T cell responses could cross-recognize the MPXV derived predicted epitope pools (**Figure 3** and **Figure S2**). In the case of CD4^+^ T cells, in both AIM and ICS assays we found a considerable cross-recognition of the MPXV megapool, and its magnitude followed similar kinetics to those observed for the Orthopox Megapool responses (**Figure 3A-B**).

**Figure 3.**
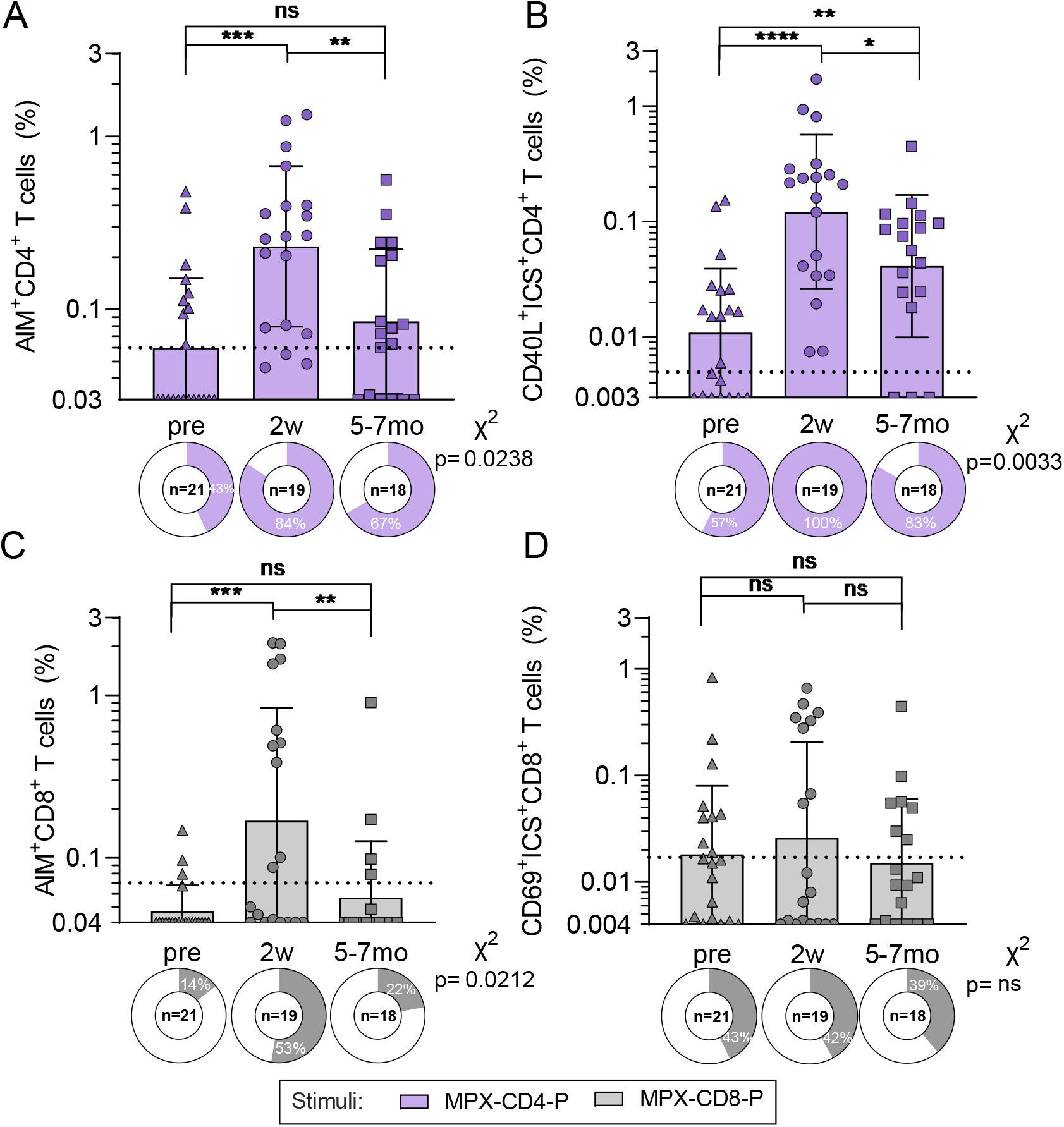
T cell responses able to cross-recognize MPXV MPs after Vaccinia vaccination. Refers to Figure S1–S2 and Supplemental Tables 1 to 4. PBMCs from each time-point were tested in AIM/ICS assays with predicted MPXV MPs, MPXV-CD4-P (purple) and 5 pools of MPXV-CD8-P1-P5 summed to get the overall MPXV-CD8-P reactivity (grey). MPXV-CD4-P specific CD4^+^ T cell reactivity are shown for (**A**) AIM and (**B**) ICS. MPXV-CD8-P specific CD8^+^ T cell reactivity are shown for (**C**) AIM and (**D**) ICS. The Y axis of each bar graph starts at the LOD and the LOS is indicated with a dotted line. Pie charts below each bar indicate the frequency of positive responders. Mann-Whitney T test was applied to each graph and p values symbols are shown when significant. * p<0.05, **p<0.01, ***p<0.001, ****p<0.0001. X^2^ test was applied to the frequencies of positivity and p values are listed on the right of each graph.

Specifically, by the AIM assay the magnitude of CD4^+^ T cell cross-reactivity peaked at 2 weeks post-vaccination and declined at 6-7 months, reaching comparable reactivity to what was observed pre-vaccination (GM±GSD; pre=0.06±2.51, 2 weeks=0.23±2.91, 5-7 months=0.08±2.63; P=0.0005 Kruskal-Wallis test; Mann-Whitney comparisons in **Figure 3A**). The frequency of CD4^+^ T cell responders significantly increased from 43% to 84% 2 weeks post-vaccination, with a decline to 67% 5-7 months post-vaccination (χ^2^ p= 0.024). Similar results were observed by ICS, although the post-vaccination decline was less pronounced and responses were still significantly higher than those observed pre-vaccination (GM±GSD; pre=0.011±3.58, 2 weeks=0.12±4.67, 5-7 months=0.04±4.13; P<0.0001 Kruskal-Wallis test, Mann-Whitney comparisons **Figure 3B**). Finally, the functionality of CD4^+^ T cells was comparable to what was observed for the OPXV-specific T cell responses (**Figure S2B** and **S2E**).

CD8^+^ T cell responses were also able to cross-recognize the MPXV pools. Reactivity peaked at 2 weeks and further declined 5-7 months post-vaccination, with comparable reactivity with the pre-vaccination by AIM (GM±GSD; pre=0.05±1.44, 2 weeks=0.17±4.93, 5-7 months=0.06±2.23; P=0.0014 Kruskal-Wallis test; χ^2^ p= 0.0212; Mann-Whitney comparisons in **Figure 3C**) and a less pronounced decline by ICS (GM±GSD; pre=0.02±4.42, 2 weeks=0.03±7.97, 5-7 months=0.02±3.96; P=0.8653 Kruskal-Wallis test; χ^2^ p= 0.9961; Mann-Whitney comparisons in **Figure 3D**). The quality of CD8^+^ T cells was comparable and driven by IFNγ production, although an increase in IL-2 was also observed (**Figure S2D** and **S2F**). Little or no reactivity was once again observed in non-vaccinated subjects for both CD4^+^ and CD8^+^, in line with the lack of exposure and absence of pre-existing cross-reactive responses (**Figure S1D-E**).

## Discussion

There is an urgent need to understand the adaptive immune responses to MPXV in the context of both natural immunity and vaccination. The present study is focused on CD4^+^ and CD8^+^ T cell responses. We defined dominant cross-reactive ORFs to generate peptide MPs to measure T cell responses, which we plan to make available to the research community. These reagents were validated by assessing T cell responses induced by Dryvax vaccination in banked PBMC samples.

Because all current vaccinations against MPXV are based on VACV/MVA, it was important to determine the sequence conservation frequency of previously defined VACV T cell epitopes and those curated in the IEDB with the MA001 strain, reference for the current outbreak. Overall, the median degree of amino acid sequence conservation was 61-67%, implying that responses induced by VACV vaccination should recognize ortholog protein sequences in the MPXV genome. McKay and collaborators reported a similar sequence conservation frequency of 70% between MPXV-2022 and the MPXV-CB (Congo Basin) strains (Ahmed et al., 2022). This observation enabled the definition of Orthopox-specific megapool reagents.

The epitope data in the IEDB was utilized to define dominant ORFs recognized in general by Orthopox specific T cell responses. This analysis revealed that the breadth of both CD4^+^ and CD8^+^ T cell immune responses is remarkable, consistent with earlier observations (Jing et al., 2008; Moutaftsi et al., 2007). Indeed, 19 ORFs were required to cover 67% of CD4^+^ T cell responses and 40 ORFs were required to cover 61% of CD8^+^ T cell responses. This is significant, but suggests that focusing on a manageable subset of the over 200 ORFs typically encoded in pox genomes will nevertheless allow the majority of T cell responses to be captured. This observation enabled the definition of MPXV-specific megapool reagents as discussed below. Furthermore, the definition of dominant ORFs provided insights into the mechanisms underlying the development of these responses showing that dominant antigens are predominantly early ORFs, enriched in structural proteins in the case of CD4^+^ T cell responses, confirming earlier studies (Moutaftsi et al., 2007). In the context of vaccination, we note that all dominant ORFs were conserved in MVA/Jynneos, and had orthologs in MPXV. This suggest that the response directed to MPXV induced by MVA should be able to mirror the response induced by the previous VACV vaccines, with known clinical efficacy against MPXV in humans.

Based on the analysis of Orthopox epitopes curated in the IEDB, and the parallel determination of MPXV orthologs of immunodominant ORFs, we developed peptide pools based on either experimentally defined Orthopox T cell epitopes or predicted T cell epitopes derived from the most immunodominant ortholog proteins of MPXV, following a similar approach to the one we previously applied in the context of SARS-CoV-2 (Grifoni et al., 2020a; Grifoni et al., 2020c). These Orthopox epitope pools were validated using banked PBMC from subjects who received the Dryvax vaccine. Early time points (2 weeks from vaccination) were associated with positive responses in 100% of the subjects for CD4^+^ T cells and 63% for CD8 T cells and decreased 5-7 months after vaccination. CD4^+^ and CD8^+^ T cell responses to the pools of predicted MPXV epitopes were similarly detected in 84% and 52% of vaccinees 2 weeks post vaccination, and also decreased 5-7 months after vaccination. To the best of our knowledge, this is the first demonstration of the use of epitope megapools to detect Orthopox specific responses in humans, and in particular the first detection of responses to MPXV-specific sequences. The use of epitope pools has the important potential advantage of obviating the use of infected cells to quantify T cell responses, which is prone to interference by pox-virus expression of immune antagonizing genes (Seet et al., 2003), and in the case of MPXV in particular, is associated with biosafety concerns. The SARS-CoV-2 peptides pools developed using a similar approach (Grifoni et al., 2020a) have been widely used in over 60 different studies to date.

The validation of the epitope pools also revealed some important new aspects of human T cell responses to VACV. Our study confirmed that CD8^+^ T cell responses are less durable than their CD4^+^ counterpart (Amara et al., 2004). At the same time, we observed that human CD4^+^ T cell responses are associated with a large fraction (up to 53%) of Granzyme B secreting antigen-specific cells. It was previously reported that the role of CD4^+^ T cells in protection from VACV and MPXV infections outweighs the contribution provided by CD8^+^ T cells in macaques (Edghill-Smith et al., 2005a; Edghill-Smith et al., 2005b). The current data suggests that in addition to their role supporting the development of antibody responses, their longevity and a cytotoxic component could also contribute to a sustained antiviral function of CD4^+^ T cells.

In conclusion, the use of available information related to VACV epitopes in conjunction with bioinformatic predictions points to specific regions that are conserved across several OPXV species, including MPXV, making them suitable for vaccine evaluation. The results also describe megapools (MPs) of peptides suitable to characterize vaccine-specific responses, and also likely to detect immune responses in the context of MPXV infection and disease.

### Limitations

The use of a cross-sectional cohort decreases the accuracy of kinetic analysis of responses, and it was not possible to exclude or confirm childhood vaccination status. Additionally, this cohort included also healthcare workers and we cannot exclude previous occupational vaccination before the Dryvax vaccination administered in relation to this specific study. This possibility is confirmed by a lower reactivity observed in a younger independently collected population when compared with the pre-vaccinated samples of our vaccinated cohort. Antigen selection for MPXV was based on studies performed considering other OPXV (mainly VACV), and we cannot therefore exclude that important antigens only recognized in the case of MPXV might have not been considered. The current analysis did not address antibody responses, which are the dominant correlate of vaccine induced protection. Future directions will include: (1) use of the megapools to characterize immune responses in acute and convalescent MPXV natural infection, (2) addressing the Th and memory phenotypes of responding T cells, and (3) comparison of responses induced by different vaccines, (4) wide dissemination of the megapools to the scientific community and (5) further optimization of these reagents.

## Acknowledgments

This project has been funded in whole or in part with Federal funds from the National Institute of Allergy and Infectious Diseases, National Institutes of Health, Department of Health and Human Services, under Contract No. 75N93019C00001 and 75N9301900065 to A.S and HHS75N93019C00076 to R.H.S. A.T. was supported by a PhD student fellowship through the Clinical and Experimental Immunology Course at the University of Genoa, Italy. All the authors declare no competing interests. LJI has filed for patent protection for various aspects of T cell epitope and vaccine design work. Please contact A.S. for aliquots of synthesized sets of peptides identified in this study. There are restrictions to the availability of the peptide reagents due to cost and limited quantity.

## Author contribution

Conceptualization: A.G., Y.Z, R.H.S and A.S.; Data curation and bioinformatic analysis, A.G., J.S. Y.Z.; Formal analysis: Y.Z. A.T., A.G.; Funding acquisition: R.H.S and A.S.; Investigation: A.T., M.R. and A.G.; Resources: J.D., P.R. supervision: G.F., A.G., R.H.S and A.S; Writing: A.G., Y.Z, R.H.S and A.S.

## Methods

### IEDB analysis of Orthopoxvirus-derived T cell epitopes

Known OPXV-derived T cell epitopes reported in the published literature, or through direct database submission, were identified by searching the IEDB at the end of May, 2022. Queries were performed broadly for the Orthopox genus, using NCBI taxonomy ID 10242, and specifying positive T cell assays. This retrieved 1076 records, from which 31 were removed because responses had not been defined in the context of either MHC class I or class II. For epitopes with responses in the context of class II, the set was further filtered to select epitopes of 12-25 residues, comporting with the canonical size of class II ligands associated with CD4^+^ T cell responses. Epitopes with class I responses were filtered to select those of 8-11 residues, canonical for class I ligands associated with CD8 T cell responses. As a result, a final set of 977 epitopes, including 318 associated with class II responses, and 659 with class I responses, was identified for subsequent analyses. About 70% of the epitope data in this set is derived from the peer-reviewed literature.

### Identification of Orthopox T cell epitope homologs in MPXV

The protein sequences of MA001 were retrieved from the BV-BRC website (https://www.bv-brc.org/view/Genome/10244.322#view_tab=overview) on May 25, 2022. In order to identify the MPXV protein region homologous to the OPXV T cell epitopes, a k-mismatch string search method was used. Conceptually, the k-mismatch string search program searches through a protein sequence file using a fixed-size sliding window and identifies all windows with a maximum of k-mismatches compared to the input epitope sequence. In identifying OPXV epitope homologs in MPXV, the search pool used included all MA001 protein sequences, while the maximum number of k mismatches was set to be the larger of 20% of the input epitope length and 1, i.e., k = max (epitope length * 0.2, 1). We additionally set up a maximum of 2 and 3 mismatches for class I and class II epitopes, respectively.

Besides finding all epitope homologs in proteins, the epitope search program also picks the best match if multiple matches were found. The best match was defined as the one with the smallest number of mismatches, and in case of ties, the one(s) with the least shift in the start coordinate compared with the input epitope.

In validating the epitope search result, two metrics were used: (1) whether the epitope hit resides in a protein homologous to the input epitope’s parent protein identified from the pairwise analysis, (2) whether the start coordinate of the epitope hit was near the start coordinate of the input epitope. In case of a start coordinate shift of 10 or more residues, the sequences of the parent proteins were aligned and then manually examined to see if the match was a false positive. For all epitopes evaluated using these criteria, only one was found to be a false positive following manual curation and was excluded from the downstream analysis.

### T cell epitope predictions

Epitope prediction was carried out using the various dominant MPXV ORFs described above (**Supplemental Table 3**). For CD4^+^ T cell epitope prediction, we applied a previously described algorithm that was developed to predict dominant HLA class II epitopes, using a median consensus percentile of prediction cutoff ≤ 20 percentile as recommended (Paul et al., 2015b). For CD8^+^ T cell epitope prediction, we selected the 12 most frequent HLA class I alleles in the worldwide population (Middleton et al., 2003, Paul et al., 2013), using a phenotypic frequency cutoff ≥ 6%. The specific alleles included were: HLA-A*01:01, HLA-A*02:01, HLA-A*03:01, HLA-A* 11:01, HLA-A*23:01, HLA-A*24:02, HLA-B*07:02, HLA-B*08:01, HLA-B*35:01, HLA-B*40:01, HLA-B*44:02, HLA-B*44:03. HLA class I binding predictions were performed using the IEDB recommended class I prediction algorithm (as recommended in June, 2022) and selecting for each allele the top 1 percentile of peptides based on the total amino acid sequences of the 40 MPXV antigens selected. This initial list of epitopes was then filtered to eliminate redundancies and nested peptides by clustering (Dhanda et al., 2018) to a single occurrence, and nested peptides were included within longer sequences, up to 12 residues in length, before assigning the multiple corresponding HLA restrictions for each region.

### Peptide synthesis and Megapool preparation

OPXV and MPXV peptides were synthesized as crude material (TC Peptide Lab, San Diego, CA), and then individually resuspended in dimethyl sulfoxide (DMSO) at a concentration of 10–20 mg/mL. Aliquots of all peptides were pooled into megapools (MP) designated as OP-CD4-E, OP-CD8-E, MPX-CD4-P, MPX-CD8-P1, MPX-CD8-P2, MPX-CD8-P3, MPX-CD8-P4, and MPX-CD8-P5. These MPs underwent a sequential lyophilization. The resulting lyocake was resuspended in DMSO at 1 mg/mL, as previously described (Grifoni et al., 2020c). All peptides and MPs are listed in **Supplemental Tables 1** and **4**.

### Cohort of VACV vaccinees to assess T cell responses

The characteristics of the study population that donated the banked PBMC utilized for the present study was described previously (Oseroff et al., 2005). Healthy male and female donors between 7 and 62 years of age that had received a vaccinia virus (Dryvax) vaccination as a prophylactic measure, either because of their potential exposure to vaccinia in a laboratory or hospital setting, or because of their enrollment into military and health worker vaccination programs, within one year of providing the blood donation. PBMCs were collected at 2 weeks (n=19) or 5-7 months after vaccination (n=18). In addition, we also utilized two additional control cohorts. For the first control cohort, Pre-vaccination (“pre”) PBMC samples (n=21) were collected from a similar cohort of healthcare donors prior to vaccination. Since some of the pre-vaccinated donors had received childhood Smallpox vaccination, a second control cohort of truly unexposed donors (n=15) was enrolled; this cohort consisted of individuals born after 1980, the official year of worldwide eradication of Smallpox and eight years after the United States stopped childhood Smallpox vaccinations. Further, this unexposed cohort had no history of occupational vaccination with Smallpox. For all donors, PBMCs were isolated from heparinized blood by gradient centrifugation with a Histopaque-1077 and cryopreserved in liquid nitrogen in 10% DMSO/FBS. Characteristics of the donor cohorts are also summarized in **Supplemental Table 2**. Institutional Review Board approval and appropriate consent were obtained for this study.

### AIM/ICS assay

We performed the combined AIM/ICS assay as previously described (Tarke et al., 2022). In brief, after thawing, 1-2×10^6^ PBMCs per well were cultured with the OPXV-or MXPV-specific peptide MPs [1 μg/mL]. An equimolar amount of DMSO was added to the cells in triplicate wells as a negative control. Phytohemagglutinin (PHA, Roche, 1 μg/mL) was used to stimulate cells as a positive control. Treated cells were incubated at 37°C in 5% CO^2^ for 22 hours before the addition of Golgi-Plug containing brefeldin A, Golgi-Stop containing monensin (BD Biosciences, San Diego, CA), and the CD137 APC antibody (2:100, Biolegend, Cat# 309810) for an additional 4-hour incubation. Then the cells underwent membrane surface staining for 30 minutes at 4°C protected from light with Fixable Viability Dye eFluor506 (1:1000, eBiosience, Cat# 65-0866-14) and the following antibodies: CD3 BUV805 (1:50, BD, Cat# 612895), CD8 BUV496 (1:50, BD, Cat# 612942), CD4 BV605 (1:100, BD, Cat# 562658), CD14 V500 (1:50, BD, Cat# 561391), CD19 V500 (1:50, BD, Cat# 561121), CD14 V500 (1:50, BD, Cat# 561391), CD69 PE (1:10, BD, Cat# 555531), CD137 APC (1:50, Biolegend, Cat# 309810), and OX40 PE-Cy7 (1:50, Biolegend, Cat# 350012). After staining, the cells were fixed with 4% paraformaldehyde (Sigma-Aldrich, St. Louis, MO), permeabilized with saponin buffer (0.5% saponin [Sigma-Aldrich, St. Louis, MO], 1% bovine serum albumin, and 0.1% sodium azide), and blocked for 15 minutes with 10% human serum (Gemini Bio-Products, Sacramento, CA) in saponin buffer. After blocking, the cells were stained intracellularly for 30 minutes at room temperature with the following antibodies: TNFα ef450(3:100, Life Tech, Cat# 48-7349-42), IFNγ FTIC (1:100, Invitrogen, Cat# 11-7319-82), IL-2 BB700 (1:25, BD, 566405), IFNγ (1:100, Invitrogen, Cat# 11-7319-82), Granzyme B Alexa700 (1:100, BD, 560213), and CD40L APC-eFluor780 (3:100, eBioscience, Cat# 47-1548-42). All samples were acquired on a ZE5 5-laser cell analyzer (Bio-Rad laboratories) and were analyzed with FlowJo software (Tree Star Inc.).

The data was analyzed to establish the Limit of Detection (LOD) and Limit of Sensitivity (LOS) based on all the DMSO-only conditions for AIM and ICS. For ICS these calculations were done on the IFNγ data. The LOD was calculated as twice the upper 95% confidence interval of the geometric mean and the LOS was calculated as two times the standard deviation from the median. Only responses with Stimulation Index (SI) > 2 were considered significant for AIM (CD4: LOS = 0.06%, SI > 2; CD8: LOS = 0.07%, SI > 2). For ICS, responses with SI>2 were considered significant for CD4^+^ (LOS = 0.006%) and responses with SI>3 for CD8^+^ (LOS = 0.017%). For AIM, the CD8^+^ T cell response to MPXV-CD8-P was calculated by summing the background subtracted, SI>2 and >LOS AIM data. For ICS, the same calculation was performed for the ICS data but considering an SI>3.

---

**Figure S1.**
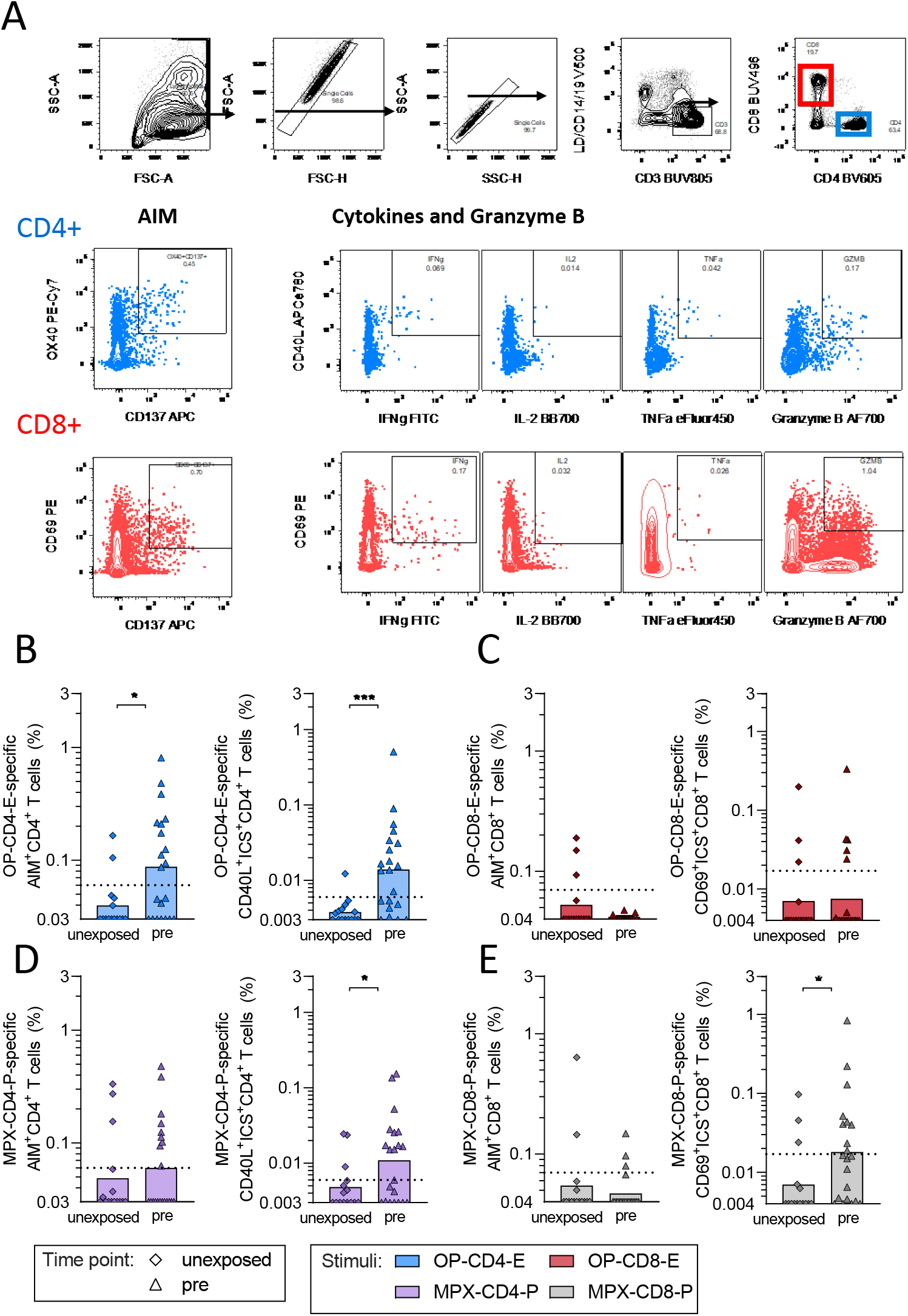
Gating strategy for AIM+ICS assay and T cell reactivity in unexposed and pre-vaccinated individuals. Related to Figures 1 and 3. (A) Gating strategy for the AIM/ICS assays used in the study. The indicated AIM or intracellular markers were used to assess OPXV or MPXV specific T cell reactivity. For ICS, the total reactivity was calculated from cells positive for IFN *γ*, IL-2, TNFα or GZMB in combination with CD40L for CD4+ or CD69 for CD8+, with the exception of single positive GZMB cells which were excluded from the CD8 sum. Unexposed individuals (n=15, diamond) and pre-vaccinated healthcare workers (n=21, triangle) were tested in AIM+ICS assays for (B) CD4+ and (C) CD8+ T cell reactivity to the OPXV megapools, OP-CD4-E (blue) and OP-CD8-E (red). The same donors were tested for (D) CD4+ and (E) CD8+ T cell reactivity for MPXV predicted peptide pools, MPX-CD4-P (purple) and MPX-CD8-P (grey) respectively. The Y axis of each bar graph starts at the LOD and the LOS is indicated with a dotted line. Mann-Whitney T test was applied to each graph and p values symbols are shown when significant. * p<0.05, ***p<0.001.

**Figure S2.**
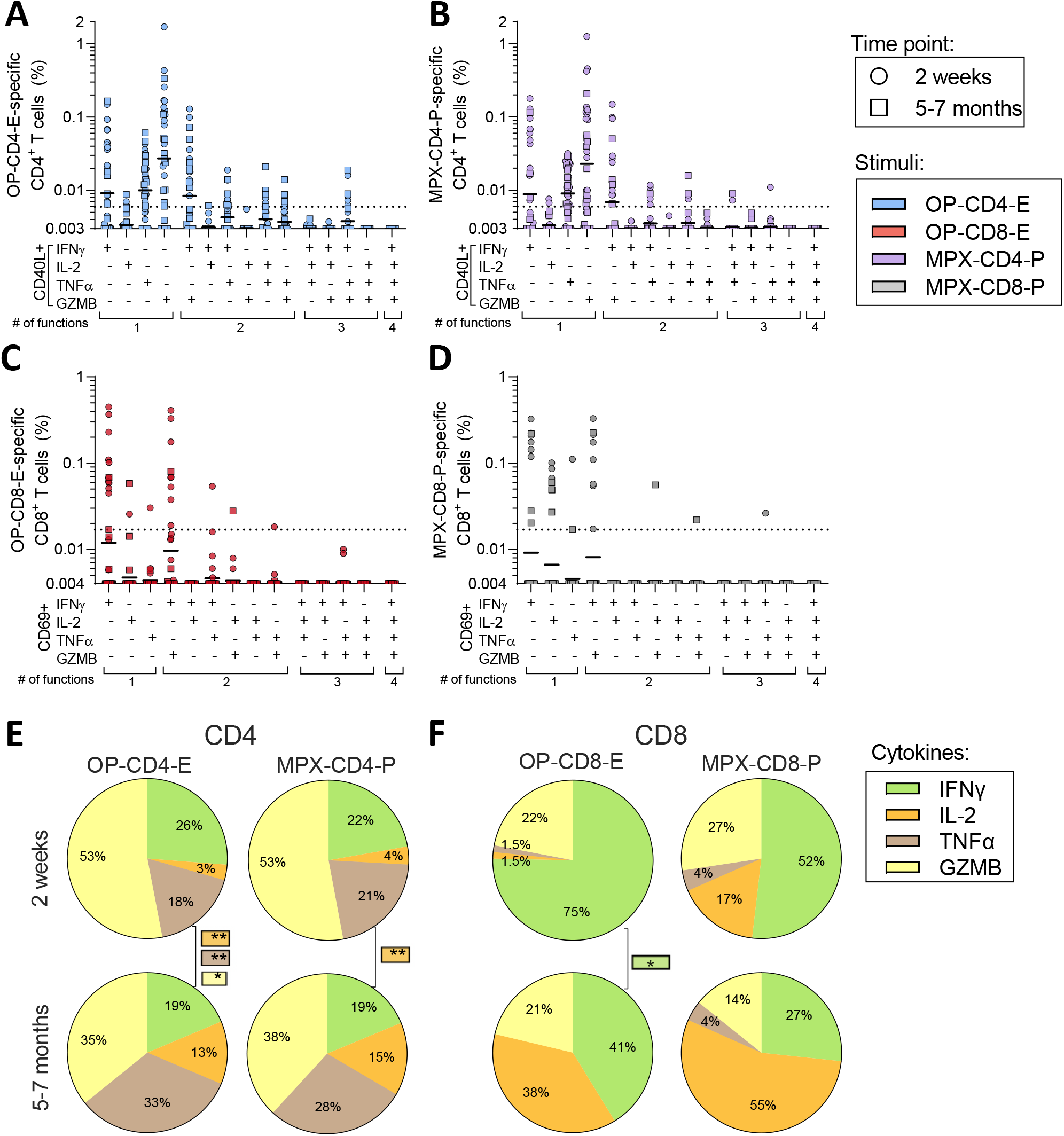
Quality of OPXV and MPXV-specific-T cell responses. Related to Figures 1 and 3. PBMCs were collected from donors two weeks after vaccination with Dryvax (n=19, circle) and five to seven months after vaccination (n=18, squares) and tested in AIM+ICS assays for T cell reactivity to OPXV and MPXV. (A – B) Boolean gating of (A) OP-CD4-E- or (B) MPX-CD4-P-specific CD4+ ICS reactivity is shown for CD40L+ CD4+ T cells in combination with IFNy, IL-2, TNFα and GZMB. (C - D) Boolean gating of (C) OP-CD8-E- or (D) MPX-CD8-P-specific CD8 ICS reactivity is shown for CD69+ CD8 T cells in combination with IFNy, IL-2, TNFα or GZMB. The Y axis of each bar graph starts at the LOD and the LOS is indicated with a dotted line. Contribution of IFNy, IL-2, TNFα and GZMB to the total of (E) CD4 or (F) CD8+ ICS responses to OP and MPX megapools at two weeks and 5-7 months post vaccination. Mann-Whitney T test was applied to each pie chart and p values symbols are shown when significant. * p<0.05, **p<0.01.

